# Signatures of Dobzhansky-Muller Incompatibilities in the Genomes of Recombinant Inbred Lines

**DOI:** 10.1101/021006

**Authors:** Maria Colomé-Tatché, Frank Johannes

**Author notes:** Corresponding authors: European Research Institute for the Biology of Ageing, University of Groningen, University Medical Center Groningen, A. Deusinglaan 1, NL-9713 AV Groningen, The Netherlands, and Groningen Bioinformatics Centre, University of Groningen, Nijenborgh 7, 9747 AG Groningen, The Netherlands,.

## Abstract

In the construction of Recombinant Inbred Lines (RILs) from two divergent inbred parents certain genotype (or epigenotype) combinations may be functionally “incompatible” when brought together in the genomes of the progeny, thus resulting in sterility or lower fertility. Natural selection against these epistatic combinations during inbreeding can change haplotype frequencies and distort linkage disequilibrium (LD) relations between loci within and across chromosomes. These LD distortions have received increased experimental attention, because they point to genomic regions that may drive Dobzhansky-Muller-type of reproductive isolation and, ultimately, speciation in the wild. Here we study the selection signatures of two-locus epistatic incompatibility models and quantify their impact on the genetic composition of the genomes of 2-way RILs obtained by selfing. We also consider the biases introduced by breeders when trying to counteract the loss of lines by selectively propagating only viable seeds. Building on our theoretical results, we develop model-based maximum likelihood (ML) tests which can be employed in pairwise genome scans for incompatibility loci using multi-locus genotype data. We illustrate this ML approach in the context of two published *A.thaliana* RIL panels. Our work lays the theoretical foundation for studying more complex systems such as RILs obtained by sibling mating and/or from multi-parental crosses.

---

Hybrids from crosses between two divergent parental lines sometimes display low fertility and phenotypic abnormalities (Presgraves 2010). These effects are often attributable to combinations of parental gentoypes (or epigenotypes) at two or more loci that are functionally incompatible when brought together into a single genome. This form of negative epistasis was originally invoked by Dobzhansky (Dobzhansky 1937) and Muller (Muller 1942) as a model for speciation. In the classical Dobzhansky-Muller (DM) model, a population splits into two sub-populations which become reproductively isolated through geographic or temporal mechanisms (i.e. prezygotically). Once separated, the two sub-populations acquire independent mutations that are incompatible upon hybridiza-tion, thus resulting in sterility or reduced fertility among offspring. This process prevents further mixing and reinforces the existing (pre-zygotic) reproductive isolation genetically (i.e. post-zygotically). Additional independent mutations accumulate over time, causing further divergence between subpopulations and ultimately speciation. Empirical examples of inter-specific genetic incompatibilities are well documented in the literature (Presgraves 2010) and have motivated extensive theoretical work in evolutionary genetics (e.g. Nei *et al.* 1983; Orr and Orr 1996; Turelli and Orr 2000; Orr and Turelli 2001; Turelli *et al.* 2001; Barton 2001; Welch 2004; Fierst and Hansen 2010; Bank *et al.* 2012). Interestingly, genetic incompatibilities with varying degrees of penetrance are often already visible in intra-specific experimental crosses of inbred laboratory strains (Corbett-Detig *et al.* 2013). The detection and functional analysis of such intra-specific incompatibilities could provide fundamental insights into the mechanisms that drive post-zygotic reproductive isolation in the wild, and thus represent a useful model for understanding the molecular basis of speciation (Bomblies and Weigel 2007).

In plants, the clearest examples of intra-specific genetic incompatibilities come from experimental crosses of *A. thaliana* (e.g. Bomblies *et al.* 2007; Bikard *et al.* 2009; Durand *et al.* 2012; Chae *et al.* 2014). Arguably the best studied case is the work of Bikard et al. (Bikard *et al.* 2009), who examined *F*_2_ progeny of selfed hybrids derived from the Columbia (Col) and the Cape Verte (Cvi) accessions. The authors found that a subset of the *F*_2_s had severely compromised fitness, and demonstrated that this fitness loss is caused by a genetic incompatibility involving a reciprocal loss of duplicate genes on chromosome (chr) 1 and chr 5 (Fig. 1A). Specifically, it was shown that Cvi carries a deletion of the gene on chr 5 and Col a non-functional version on chr 1, both of which act recessively. Hence, *F*_2_ individuals with the recessive epistatic combination Col *|*Col (chr 1) and Cvi *|*Cvi (chr 5) are (nearly) embryonic lethal. Interestingly, the genomic regions that are implicated in this epistatic incompatibility were first identified in a densely genotyped population of Recombinant Inbred Lines (RIL) derived from the Col and Cvi accessions: at generation *F*_6_ of inbreeding, the authors noted strong long-range linkage disequilibrium (LD) between markers on chr 1 and 5 (Fig. S1A) (Simon *et al.* 2008). Combinations of the Col Col marker genotype on chr 1 and the Cvi Cvi marker genotype on chr 5 were completely absent, suggesting that these epistatic combinations were subject to intense selection during inbreeding. Similar long-range LD patterns were identified in another RIL population originating from Shahdara (Sha) and Col accessions, and involved epistatic interactions between a locus on chr 4 and chr 5 (Fig. S1B). The two loci were subsequently fine-mapped, and functional studies revealed that this epistatic incompatibility is due to stable epigenetic silencing of a paralogue (Fig. 1B) (Durand *et al.* 2012). This latter finding illustrates that beside genetic factors also epigenetic factors can cause intra-specific incompatibilities in plants.

**Figure 1 (A).**
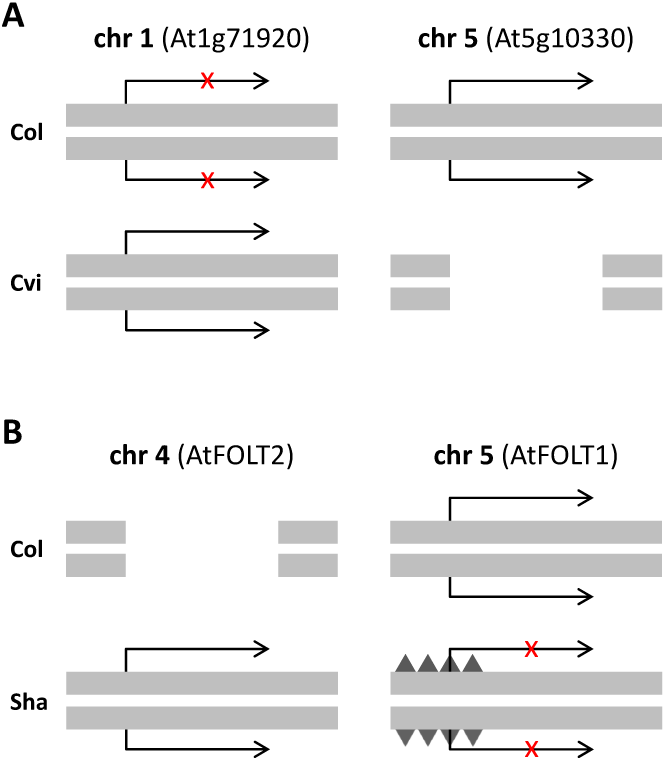
Example genetic incompatibility in a cross between *A. thaliana* accessions Col and Cvi. Locus At1g71920 on chr 1 is expressed in Cvi but not in Col, while homologous locus At5g10330 on chr 5 is expressed in Col but deleted in Cvi. (**B**) Example genetic incompatibility in a cross between *A. thaliana* accessions Col and Sha. Locus AtFOLT2 on chr 4 is expressed in Sha but deleted in Col, while homologous locus AtFOLT1 on chr 5 is expressed in Col but epigenetically silenced in Sha through DNA methylation (black triangles).

Shortor long-range LD distortions between loci on the same or on different chromosomes is a common feature of RILs genomes. From the standpoint of complex trait analysis, such distortions are typically undesirable because they affect the resolution and power of quantitative trait locus (QTL) mapping. On the other hand, a systematic analysis of LD distortion patterns can provide insights into the epistatic architecture underlying genetic incompatibilities and yield targets for experimental follow-up. A decisive contribution to such efforts is a theoretical analysis of different incompatibility models and their selection signatures in the genomes of RILs. Most of the theoretical work devoted to understanding the genomes of RILs has ignored the role of selection (e.g. Haldane and Wadington 1931; Broman 2005; Martin and Hospital 2006; Teuscher and Broman 2007; Johannes and Colomé-Tatché 2011; Martin and Hospital 2011; Broman 2012; Zheng *et al.* 2015). The exception is the early work by Haldane (Haldane 1956), Reeve (Reeve 1955) and Hyman and Mather (Hayman and Mather 1953), who examined cases of selection against homozygotes at a single locus, and described the changes in genotype frequencies as a function of inbreeding and selection. However, these earlier theoretical results are of limited use for understanding the selection signatures of DM-type genetic incompatibilities as the latter require multi-locus models.

Here we provide the first theoretical analysis of two-locus incompatibility models in the context of RIL construction. We consider three variants of the classical DM-model (the dominance epistasis, the recessive epistasis, and the dominance-recessive epistasis model) and quantify their respective effects on short-and long-range LD patterns as a function of inbreeding, fitness and recombination. We also give theoretical expressions for the total number of lines that are expected to be lost under different incompatibility scenarios. Building on these results, we present model-based maximum likelihood (ML) tests which can be used for the detection of incompatible loci from multi-locus genotype data collected at any inbreeding generation. We apply this ML method to two published *A. thaliana* RIL panels. Our work lays the theoretical foundation for studying more complex systems such as RILs obtained by sibling mating and/or from multi-parental crosses.

## Overview of genetic incompatibility models

The simplest form of epistatic incompatibility involves the interaction between only two loci, say *L*_1_ and *L*_2_. Consider two divergent inbred lines with diplotypes (i.e. two-point genotypes) 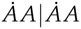 and 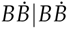, where the “dot” superscript denotes a nonfunctional (i.e. mutant) allele. We use the notation *IK| J L* to distinguish genotypes *I| J* and *K |L* at the first (*L*_1_) and second (*L*_2_) locus, respectively, from haplotypes *IK* and *JL* on each of the two homologous chromosomes (Table 1). Hence, inbred line 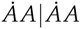 is homozygous for two mutant alleles at the first locus and homozygous wild type at the second locus, while inbred line 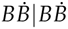 is homozygous mutant at the second locus and homozygous wild-type at the first locus. There are three basic models of two-locus epistatic incompatibility, the dominance epistasis model (*M*_1_), the recessive epistasis model (*M*_2_), and the dominance-recessive epistasis model (*M*_3_). These models are summarized in (Table 2) and are further detailed below.

**Table 1.**
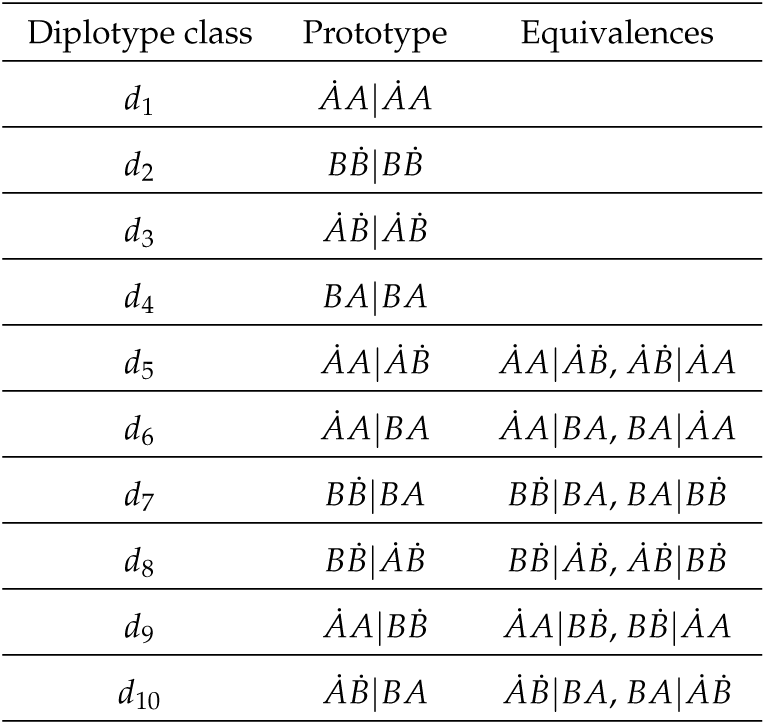
List of the 16 diplotypes arising from the 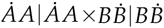 cross, where A and B denote wild-type alleles and 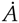 and 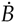 denote non-functional (mutant) alleles. Ignoring hap-lotype order the 16 diplotypes can be grouped into only 10 different classes

**Dominance epistasis model (***M*_1_): In the classical DMmodel, individuals with diplotypes 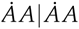 and 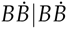 are fully viable, but their *F*_1_ hybrid progeny 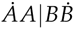 is sterile or shows reduced fertility. The reduced fitness of the hybrid is the result of dominance interactions of loci *L*_1_ and *L*_2_, meaning that allele 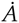 is dominant over *B* at *L*_1_, while allele 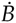 is dominant over *A* at *L*_2_. When the loss of fertility is not fully penetrant, *F*_1_ hybrids can be crossed (or selfed) to obtain a *F*_2_ population. Due to recombination and/or independent segregation of alleles at loci *L*_1_ and *L*_2_, there are 16 possible diplotypes in the *F*_2_ (Table 1). One can assume that double heterozygote *F*_2_ individuals (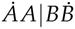) experience the same loss of fitness as in the *F*_1_. However, due to the dominance interactions, there are additional diplotypes in the *F*_2_ or in subsequent generations that are phenotypically equivalent to 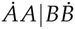, and will therefore be subject to the same, or similar, fitness loss. These diplotypes, with their corresponding fitness parameters *w*_*j*_, are summarized in Table 2.

**Table 2.**
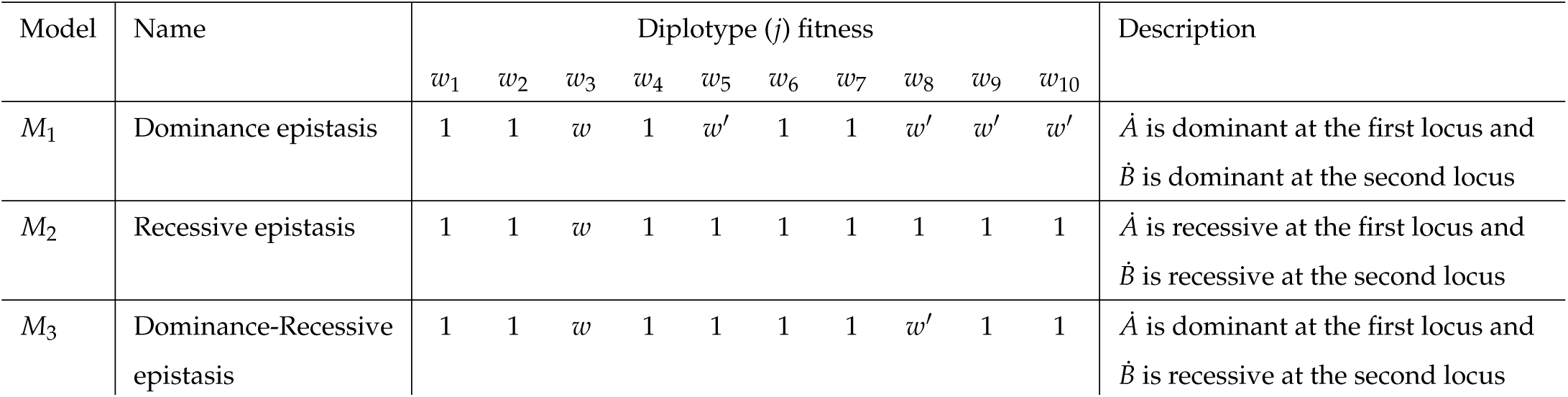
Overview of the three incompatibility models *M*_1_, *M*_2_ and *M*_3_. Shown are the fitness parameters assigned to each diplotype *j* (Table 1) with *w*, *w′ <* 1.

**Recessive epistasis model (***M*_2_): A basic requirement of the classical DM model is that the incompatibility appears in *F*_1_ hybrids. This may not always be the case. A less stringent version of the DM model is the recessive epistasis model. In this model allele *A*? is recessive to *B* at the first locus and allele *B*? is recessive to *A* at the second locus. This leads to selection against 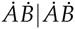 individuals, which do not appear in the *F*_1_ population but only at later breeding generations at low frequency (Table 2).

**Dominance-recessive epistasis model (***M*_3_): A combination of the dominance and the recessive epistasis model is the dominance-recessive epistasis model. In this model, allele 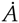 is dominant over *B* at the first locus and 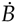 is recessive to *A* at the second locus. Selection is against individuals with diplotype 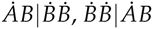 and 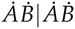 (Tables 1 and 2). Similar to the recessive epistasis case, this model implies that incompatibility does not appear in *F*_1_ individuals but only at later breeding generations. The reciprocal model where allele 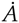 is recessive to *B* at the first locus and 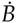 is dominant over *A* at the second locus is equivalent and can be obtained by considering the symmetries *A ↔ B* and *L*_1_ *↔ L*_2_.

Of course, the above three incompatibility models are just as valid had we assumed that the two inbred lines are instead 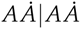 and 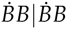, meaning that the mutant allele 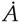 is at the second locus and mutant allele 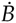 at the first locus. Various degrees of semi-dominance are taken into account by attributing different fitness parameters to deleterious diplotype (Table 2). In the following section we develop the necessary analytical framework to quantify the population-level consequences of these three incompatibility models during RIL construction. Readers who are primarily interested in the biological insights may skip directly to the Results Section.

## Theory

### Markov chain model

Consider the construction of a 2-way RIL by selfing starting from a *F*_2_ base-population. There are 16 possible diplotypes in the *F*_2_. Ignoring haplotype order, these can be grouped into 10 diplotype classes (Table 1). Individuals from the *F*_2_ (time *t* = 1) are chosen to initiate an inbreeding process by repeated selfing for many generations to obtain a final population of RILs. The inbreeding process can be modeled as an absorbing finite Markov chain, where the states of the chain are the different diplotypes *{d*_1_, …, *d*_10_ *}*(Table 1). Assume that *χ*_*t*_ denotes the diplotype state of an individual at generation *t*.

Then *{χ_t_ }* forms a Markov chain, i.e., *χ*_*t*_ + 1 is independent of *χ*_0_, *χ*_1_, …, *χ*_*t-*1_ given *χ*_*t*_. We define the transition probability *T*_*ij*_ = Pr(*χ*_*t*+1_ = *d*_*j*_ *|χ*_*t*_ = *d*_*i*_) as a function of both *r* and *w*_*j*_, where *r* (0 *≤ r ≤* 0.5) is the recombination rate at meiosis, and *w*_*j*_ (0 *≤ w_j_ <* 1) is the fitness corresponding to diplotype *j*. The transition matrix *T* is the collection of transition probabilities from one diplotype to another in one generation of inbreeding. For notational simplicity we will omit the “dot” superscript in the following, and implicitly keep track of the origin of the non-functional alleles. The general form of *T* is shown in Appendix A. Following Reeve (1955), we augment the Markov chain with a pseudo-state “lost”, which accounts for the loss of diplotypes as a result of differential survival. The column corresponding to the “lost” state in the new transition matrix *T\** is given by 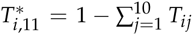 for each line *i* = (1, *…*, 10), and by 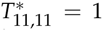 for line 11. This addition ensures that the rows of the new transition matrix *T\** sum to unity. The initial 1×11 row vector of state probabilities is

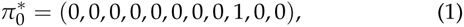

which corresponds to the hybrid diplotype *AA |BB* at *F*_1_ (time *t* = 0) where there are no other diplotypes and no selection unless *w*_9_ = 0. Hence, the state probabilities at any generation *t* of inbreeding can be obtained from the general formula

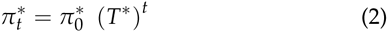

where

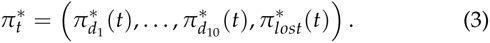

Note that the elements of 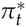 are functions of *r* and the fitness *w*_*j*_. Since only the surviving lines are of interest, one may drop the “lost” state and work instead with the reduced 10×10 submatrix of survivors, *T*, and the reduced 1×10 state vector *π*_*t*_ (Reeve 1955). This leads to the recursion

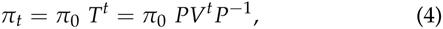

where *P* is the eigenvector of *T* and *V* is a diagonal matrix of the distinct eigenvalues of *T*. We obtain the relative diplotype proportions of surviving lines by normalizing the diplotype proportions at any generation *t* of inbreeding by the mean fitness in the population at time *t*, 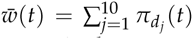 Let us define the normalized diplotype frequencies by

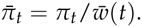

Using equation 4 we derive analytical expressions for the diplotype probabilities at any inbreeding generation (Wolfram Research Inc. 2015). For models *M*_1_, *M*_2_, *M*_3_ and for the case without selection (model *M*_0_), we list the nonnormalized diplotype probabilities at *F*_∞_ in Appendix B and D, and those for intermediate generations in Appendix C and E. The expected proportion of lost lines (*lost*) can be easily calculated from these non-normalized diplotype probabilities using

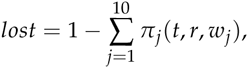

which shows that the proportion of lost lines depends on the inbreeding generation *t*, the fitness *w*_*j*_ and the meiotic recombination rate *r* between the two incompatible loci.

### Breeder Bias

In practical situations, the breeder would want to keep as many lines as possible, and therefore tries to counteract the loss of lines by implementing what may be called “biased single seed descent” (BSSD) (Fig. S2). That is, rather than selecting only one seed at random to propagate a given line to the next generation, the breeder plants many seeds from one line and chooses one that appears viable (Fig. S2). This is equivalent to arguing that the breeder will not propagate a lost line. This correction process can be modeled by normalizing each row element (*ij*) of *T* by the row total:

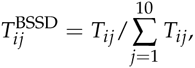

which has the effect that no lines are actually lost at intermediate generations or at *F*_∞_. The only exception is when there is complete lethality (i.e. *w*_*j*_ = 0). In this case, lines that have become fixed for a given incompatible homozygous diplotype will not produce any viable seed at all, thus leaving no alternative seeds to choose from. Although it is possible to find closed form solutions for these re-normalized diplotype probabilities, these expressions have no easy form and are therefore omitted.

### Time-dependent Linkage disequilibrium (LD)

Changes in diplotype frequencies alter haplotype proportions in the population. As we will see, all incompatibility models result in a relative gain in non-recombinant diplotypes; or stated alternatively, in a loss of diplotypes carrying recombinant haplotypes. These haplotype distortions lead to increased linkage disequilibrium (LD) within chromosomes (i.e. short-range LD) and also between chromosomes (i.e. long-range LD). To calculate LD between loci *L*_1_ and *L*_2_ we first obtain the haplotype probabilities for any time *t* as follows:

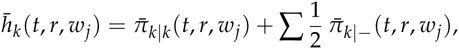

where *k ∈ {AA*, *BB*, *AB*, *BA}* and is any haplotype but *k*. Analytical expression for these haplotype probabilities for models *M*_1_, *M*_2_, *M*_3_ at generation *F*_∞_ can be found in Appendix D, and those for intermediate generations in Appendix E. As a reference we also provide the results for the case without selection (*M*_0_) in Appendix B and C (at generations *F*_∞_ and at intermediate generations, respectively). For the case of breeder bias analytical solutions are possible but have no easy form and are therefore omitted. Using these haplotype probabilities, we define the random variables *y*_1*k*_ and *y*_2*k*_ which take values 1 or 1 according to whether locus 1 or 2 on haplotype *k*, respectively, carry alleles *A* or *B*. A time-dependent measure of LD can be obtained by calculating:

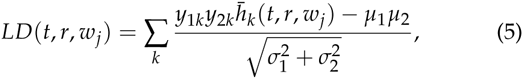

where *k ∈ {AA*, *BB*, *AB*, *BA}* and *µ*_1_, *µ*_1_ and 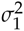, 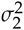 are the means and variances of *y*_1_ and *y*_2_, respectively.

### Maximum Likelihood estimation

The analytical expressions for the diplotype probabilities (Appendix E) can be employed in a Maximum Likelihood procedure for the analysis of multi-locus RIL genotype data at any generation of inbreeding. This procedure provides a method for estimating the most likely incompatibility model to have generated the data as well as the fitness coefficients corresponding to the different diplotypes.

Consider a sample of *N* RILs collected at any inbreeding generation *t*, with one random sibling representing each line. Let *Y*_*j*_ (*j* = 1, …, 10) be a random variable denoting the number of lines with diplotype *d*_*j*_ (or its equivalent class) at loci *L*_1_ and *L*_2_. Since the lines are independent, the probability mass function of the observations *y*_1_, …, *y*_10_ is given by a multinomial distribution:

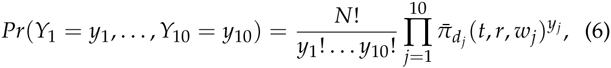

where *y*_1_ +… + *y*_10_ = *N*. Ignoring constant terms, we write the log-likelihood function (*l′*) for a given incompatibility model *M*_*i*_ and a fixed recombination fraction *r* as:

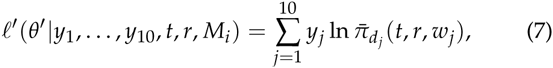

where *θ′* are the unknown fitness values. Maximization of (7) yields estimates of the fitness as well as the likelihood value of a given incompatibility model. Competing incompatibility models can be compared using standard model comparison criteria. With multi-locus marker data, the above ML approach can be used to identify loci with epistatic incompatibilities in pair-wise genome scans. It is important to point out that the meiotic recombination rate *r* needs to be fixed at its expected value during ML estimation. This is necessary because the fitness parameters are statistically confounded with *r*, so that unique solutions are not possible. However, the expected value of the meiotic recombination rate between two markers on the same chromosome is typically unknown. The only situation where *r* is known is when the two markers are on different chromosomes, in which case *r* = 0.5. The ML procedure is therefore only suited for making interference regarding long-range LD. This limitation is inherent to the problem of simultaneously estimating *r* and *w*_*j*_ and cannot be easily bypassed, irregardless of the inference method used.

## Results

The following section highlights several important biological insights that may be of practical relevance for experimentalists working on genetic incompatibility or with populations of RILs in general. Throughout we will present results for generation *F*_8_ (as this is a typical reference generation in the construction of RILs by selfing) and generation *F*_∞_ (as this is the theoretical limit), and for the three incompatibility models (*M*_1_, *M*_2_, *M*_3_) and the model without selection (*M*_0_). Results for any other inbreeding generation can be directly extracted from the analytical formulas presented in the Theory Section and the Appendix. To simplify discussion we will consider the special case *w′* = *w* (i.e. no partial-dominance).

### Genetic incompatibility leads to a loss of lines during inbreeding

The most obvious consequence of genetic incompatibility is that selection against certain diplotypes leads to the eventual loss of lines during inbreeding. The magnitude and rate of this loss depends on the mode of incompatibility (i.e. models *M*_1_, *M*_2_ and *M*_3_), the meiotic recombination rate (*r*) as well as the fitness *w*. To illustrate this, we plot the expected proportion of lost lines for two different values of *r* (0.05, 0.5) against *w* at generation *F*_8_ and *F*_∞_ (Fig. 2A).

**Figure 2.**
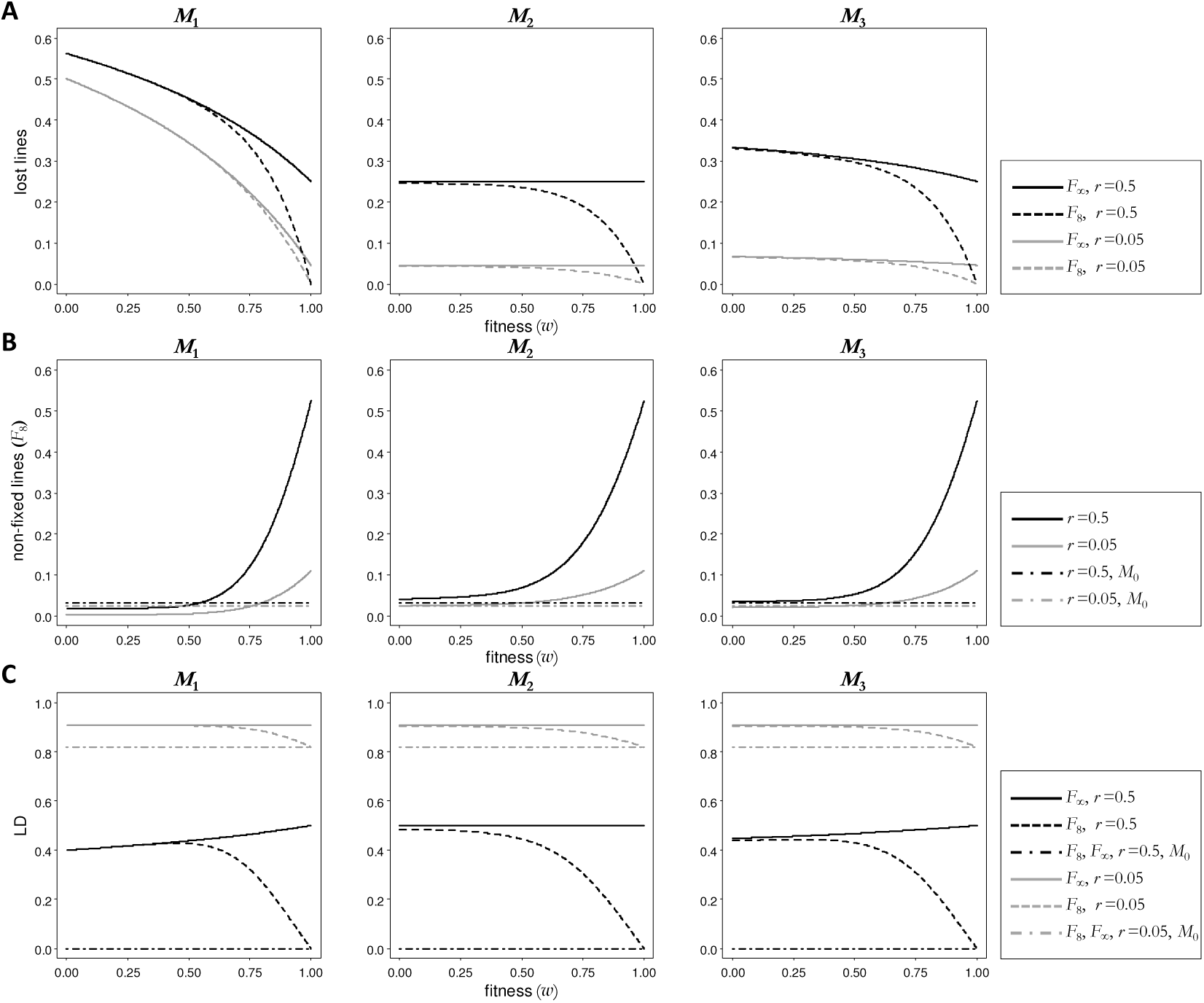
Single seed descent (SSD) results: (A) Proportion of lost lines versus fitness of the incompatible diplotypes in the SSD model. For *w* = 1 the proportion of lost lines is 0, while for w = 0 in M1 the proportion of lost lines is 1. (B) Proportion of diplotypes that have not yet reached fixation at generation F8 versus fitness for the SSD model. For *w* = 1 the proportion of non-fixed lines in *M*_1_, *M*_2_ and *M*_3_ is the same as in M0. (C) Linkage disequilibrium versus fitness for the SSD model. For *w* = 1 the linkage disequilibrium in *M*_1_, *M*_2_ and *M*_3_ is the same as in *M*_0_.

The loss of lines is most severe for the dominance-epistasis model (*M*_1_). This is because the number of different diplotypes that are selected against is largest under this model (Table 2). As the fitness of the incompatible diplotype approaches zero (*w→* 0) more than 50% of the lines are expected to be lost by generation *F*_∞_, and this percentage is not much influenced by *r*.

It is perhaps not surprising that the recessive-epistasis model (*M*_2_) is the most benign, with the loss of lines never exceeding 25% as selection acts exclusively against the genetically fixed recombinant diplotype *AB |AB*. Hence, the loss of lines at generation *F*_∞_ depends only on *r* but not *w*. With larger *r* more lines acquire recombinant haplotype *AB* during inbreeding and this haplotype can go on to fixation. By generation *F*_∞_ all *AB AB* lines will have been purged from the population. Hence, given sufficient time this processes does not depend on the selection intensity, but does require that *w* < 1.

For low fitness (*w <* 0.5) selection is generally quite efficient such that the proportion of lost lines converges rapidly to its limiting value at *F*_∞_. However, for *w* 2: 0.5 the proportion of lost lines at generation *F*_8_ differs from what is expected at generation *F*_∞_, and this feature is common to all models. Another common feature of all three models is that the loss of lines is positively related to the recombination rate between the two incompatible loci. This is because selection acts primarily against recombinant diplotypes in all models (Table 2), so that the loss of lines is expected to be most severe when the incompatibility is due to unlinked loci.

Differential survival of lines during inbreeding has other, less obvious, population-level consequences: It affects genotype and haplotype frequencies, which in turn can distort LD patterns in the genomes of RILs. We will discuss these effects in the subsequent sections.

### Genetic incompatibility changes genotype frequencies beyond fixation

In the construction of RILs by selfing inbreeding is usually not taken further than generation *F*_8_ as the lines are considered nearly fixed at that point. Indeed, in the absence of selection (*M*_0_) the *F*_8_ diplotype frequencies are close to their theoretical limit (*F*_∞_), with only about 3% of the lines still awaiting fixation. This situation is drastically different when genetic incompatibilities are present in the form of models *M*_1_, *M*_2_ and *M*_3_. In this case, certain diplotypes, many of which are already genetically fixed such as *AB| AB*, are under persistent selection and thus continue to change the relative genotype frequencies among RILs, even at very advanced inbreeding generations. To illustrate this we plot the combined frequency of all diplotypes that are still subject to change after generation *F*_8_ (Fig. 2B). For *w* 2: 0.75 all three incompatibility models show a higher frequency of changing diplotypes compared to the case without selection. One major reason for this is that selection against specific recombinant diplotypes (e.g. *AB |AB*) persists for much longer than the time it takes to generate them through recombination and fixation. This effect is clearest when the two incompatible loci are unlinked (*r* = 0.5).

With decreasing fitness the three incompatibility models begin to differ in subtle ways: for *w*,:S 0.5, the frequency of changing diplotypes after generation *F*_8_ is actually smaller for model *M*_1_ than it is for the case without selection (*M*_0_), for example for *w* = 0.3 and *r* = 0.05, the frequency of changing diplotypes after generation *F*_8_ in *M*_1_ is 0.35%, compared to 3.1% in *M*_0_. This can be attributed to the fact that many genetically un-fixed diplotypes (e.g. *AB AA*) are purged at a faster rate than the rate at which they become fixed. A similar, albeit less drastic, situation occurs in model *M*_3_ but requires much stronger selection pressures and is dependent on *r* (Fig. 2B). By contrast, in model *M*_2_ the frequency of changing diplotypes never drops below to that without selection. This is due to selection being restricted to the recombinant diplotype *AB AB*, so that fixation for this diplotype needs to occur first before it can get purged from the population.

Taken together, these results raise important practical considerations: They imply that RILs that segregate incompatible genotypes cannot be viewed as an ‘eternal’ genetic resource, as their genotype frequencies continue to change upon further propagation, particularly under weak selection. With plants this can be partly bypassed by stocking seeds from a reference generation which is then distributed to the community for phenotyping experiments. However, with RILs derived by sibling mating, lines can only be maintained by continued crossing. Experimental results obtained with genetic material from different inbreeding generations may therefore not be comparable.

### Genetic incompatibility increases LD within and across chromosomes

It is intuitively obvious that selection against certain diplotypes during inbreeding indirectly affects haplotype frequencies. Changes in the relative frequency of recombinant haplotypes distort LD relations between loci within or across RIL chromosomes. To visualize this, we plot LD against *w* for different values of the meiotic recombination rate, *r* (0.05 and 0.5), for generations *F*_8_ and *F*_∞_ (Fig. 2C). Probably the most important observation is the strong induction of long-range LD between genetically unlinked loci (*r* = 0.5) for all incompatibility models. Indeed, at generation *F*_∞_ the genotypes at the two incompatible loci are expected to be correlated in the order of 0.5, whereas they are expected to be uncorrelated in the absence of selection. For *w <* 0.5, LD(*F*_8_) ≈ LD(*F*_∞_), meaning that long-range LD rapidly reaches its maximum value with time. However, for *w* 2: 0.5 long-range LD at generation *F*_8_ and generation *F*_∞_ start to diverge substantially: long-range LD continues to increase beyond generation *F*_8_ as the relative frequency of recombinant haplotypes slowly decreases as a result of differential survival of lines. LD within chromosomes (*r <* 0.5) is of course already high due to gametic linkage, and scales with the genetic distance between the two incompatible loci. In this case, selection will reinforce LD even further. This leads to significant (local) genetic map contractions in the genomes of RILs.

One counter-intuitive observation in the LD patterns for models *M*_1_ and *M*_3_ is the slight increase in LD at generation *F*_∞_ as a function of fitness. To understand this it is necessary to discuss the fate of haplotypes during inbreeding under these two models. In both cases the proportion of recombinants depends on the fitness *w*, and both models show that low fitness values will lead to a higher proportion of recombinant diplotypes compared to higher fitness values (Fig. 3). However, recall that selection in both incompatibility models is against several diplotypes (Table 2), many of which carry the non-recombinant parental haplotypes *AA* or *BB*. Hence, with strong selection (low fitness) more lines are lost, but among the survivors there is an overrepresentation of diplotypes carrying recombinant haplotypes. By contrast, with lower selection (higher fitness) there are more surviving lines, but among these there is a higher proportion of parental non-recombinant haplotypes.

**Figure 3.**
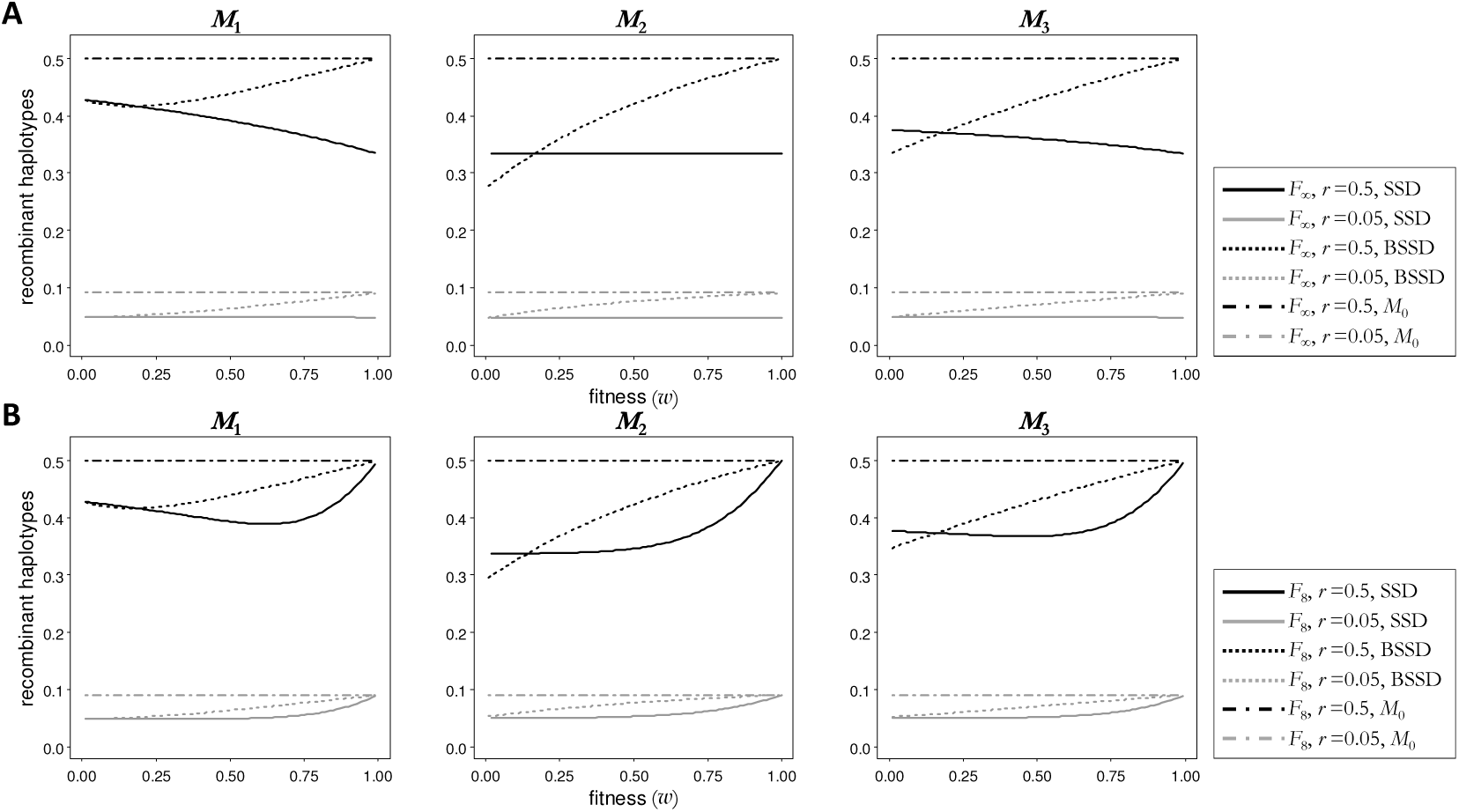
Recombinant haplotypes: Proportion of recombinant haplotypes versus fitness of the incompatible diplotypes at generation *F*_∞_ (**A**) and at generation *F*8 (**B**), for the single seed descent (SSD) and the biased single seed descent (BSSD) models. For *w* = 1 the proportion of recombinant haplotypes in *M*_1_, *M*_2_ and *M*_3_ is the same as in *M*_0_, while for *w* = 0 in *M*_1_ with SSD the proportion of recombinant haplotypes is 0.

### Preventing the loss of lines introduces additional biases

It seems sensible that many of the adverse affects of genetic incompatibility could be bypassed by preventing the loss of lines in the first place. However, preventing the loss of lines through counter-selection (BSSD, Fig. S2) does not imply that the diplotype frequencies are also corrected as if no selection had occurred. Selection against incompatible diplotypes persists, but the breeder chooses to propagate a compatible individual instead of losing a line by trying to propagate an incompatible one (Fig. S2). In this way the breeder introduces unexpected biases into the inbreeding dynamics, particularly with regards to haplotype frequencies and LD patterns. This is clearly illustrated in Figure 3, where we plot the proportion of recombinant haplotypes among surviving lines. In general, we find that BSSD leads to higher proportion of recombinant haplotypes than in the case of standard single seed descend (SSD). However, these proportions are nowhere close to what would be expected in the absence of genetic incompatibility. The most unexpected observation is that for unlinked loci, when *w*,:S 0.2, BSSD can actually produce a lower proportion of recombinant individuals among surviving lines. This means that even though more lines have been salvaged, the proportion of recombinant haplotypes in the final RILs is even lower than among surviving lines without breeder bias. The trade-off between the number of surviving lines and the proportion of recombinant individuals is important for complex trait mapping analysis where not only the sample size but also the proportion of recombinants are key determinants of mapping resolution. Making informed decisions regarding the use of BSSD is difficult, as the presence and/or severity of genetic incompatibilities is usually unknown prior to RIL construction. Be it as it may, the important observation about BSSD is that it will lead to another set of biases in genomes of RILs (Fig. 3 and Fig. S3). Breeders should be aware of these biases.

### Application to RIL genotype data

Simon et al. (2008) presented genetic maps of two *A. thaliana* RIL populations derived from crosses between Cvi×Col and Sha×Col accessions. Their analysis of the genotype data revealed several cases of long-range LD between pairs of markers on different chromosomes (Fig. S1). In the Cvi×Col cross, long-range LD was detected between markers on chr 1 and 5 and between markers on chr 1 and 3. In the Sha×Col cross, long-range LD was detected between markers on chr 4 and 5. The authors suggested that these LD patterns are the results of intense epistatic selection against certain parental genotype combinations during inbreeding. We reanalyzed the genotype data from both RIL crosses using our ML approach (Eq. 7). We focused on the two significant LD patterns originally described by Simon *et al.* (2008) and for which later experimental follow-up work established the precise mode and molecular basis of the incompatibilities (Fig. 1). In each case, we performed ML estimation using our three incompatibility models (*M*_1_, *M*_2_, *M*_3_) with and without breeder bias and, when appropriate, considered the symmetries *A ↔ B*, *L*_1_*↔L*_2_ and 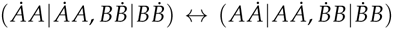 (Table S1 and S2). Our goal was to infer the most likely incompatibility model to have generated the observed genotype data, and to obtain estimates of the fitness values.

***Cvi×Col cross:*** In their analysis of the Cvi×Col cross, Simon et al. (2008) noted that individual RILs that carried the Col *|*Col genotype at a marker on chr 1 were much less likely to carry the Cvi *|*Cvi genotype at a marker on chr 5, although these loci were physically unlinked. In an impressive follow-up study (Bikard *et al.* 2009) it was later demonstrated that the chr 1 and chr 5 incompatibility was due to a reciprocal loss of a duplicated gene (Fig. 1A). Specifically, it was shown that Cvi carried a deletion of the gene on chr 5 and Col a non-functional version of it on chr 1, both of which acted recessively. *F*_2_ individuals with the recessive epistatic combination Col *|*Col (chr 1) and Cvi *|*Cvi (chr 5) were found to be (nearly) embryonic lethal. Consistent with their follow-up results in the *F*_2_s, application of our ML approach to the original RIL genotype data correctly identified the recessive epistatic incompatibility model (Model *M*_2_) as the most likely, with non-functional alleles at chr 1 for Col and at chr 5 for Cvi (Table S1). In addition, we estimated that epistatic selection against the double recessive during inbreeding was substantial (fitness *w* = 0.3234) (Table S1), which is in line with the (near) embryonic lethality observed among the *F*_2_s.

***Sha×Col cross:*** The genetic incompatibility in the Sha×Col cross is more complex: Simon et al. (Simon *et al.* 2008) observed that the combination Col *|*Col in chr 4 and Sha Sha on chr 5 was nearly absent in the RILs. Molecular analysis of the two interacting genomic regions (Durand *et al.* 2012) revealed that Sha carries a duplicated gene on chr 4, which epigenetically silences its original copy on chr 5 in *trans*. Silencing is most likely achieved via the generation of small interfering RNA (siRNA) that promote methylation at homologous sequences. Adding to this complexity, the authors showed that Sha has an active copy of the gene in chr 4, where no homologous gene exists for Col, while Col has an active copy of the gene in chr 5, where this copy is epigenetically silenced in Sha. Application of our ML approach to the genotype data revealed that the chr 4 and 5 incompatibility is most consistent with a partial dominance model (Model *M*_3_), with strong selection against individuals with genotypes Col Col on chr 4 and Sha Sha on chr 5 (*w* = 0.1250), and weak selection against individuals with genotypes Col Sha on chr 4 and Sha Sha on chr 5 (*w′* = 0.7355) (Table S2). These rather low fitness values are consistent with the authors observation that incompatible individuals experienced a reduction in seed yield of about 80%-90%. Interestingly, our ML analysis also detected evidence for breeder bias in these data, indicating that the authors made concerted efforts to counter-act the loss of lines during RIL construction, which is consistent with the breeding strategy described by the authors (Durand *et al.* 2012). Indeed, we estimate that without counterselection, approximately 30% of the lines would have been lost.

The predominance of recessive or (partial) dominance epistasis as a source of genetic incompatibilities in the Cvi×Col and the Sha×Col cross makes sense considering that other incompatibility effects such as those associated with full dominance epistasis would have led to an initial loss of *F*_1_ individuals, which may have prevented the construction of these RILs in the first place. We therefore suspect that the detection of long-range LD in multi-locus RIL genotype data will most often be traceable to recessive or partial dominance epistasis, or else to dominance epistasis in combination with weak selection.

## Discussion

Recombinant Inbred Lines (RILs) are a popular tool for studying the genetic basis of complex traits. Many populations of RILs have been created in variety of organisms. The genotypic properties of RILs often diverge drastically from what is expected from theory. Widespread allele frequency distortions and unexpected long-range LD patters are common. Such features are often the result of differential survival (or fertility) of certain combinations of parental genotypes during inbreeding. This is perhaps nowhere clearer than in the genomes of recently created 8-way RILs in the mouse, which were derived from eight different inbred parental strains (Collaborative Cross Consortium 2012). The construction of these RILs has been severely hampered by high lethality and infertility rates during inbreeding. Genotyping data at intermediate generations showed that certain parental genotypes were nearly absent in some genomic regions, and surviving lines displayed complex long-range LD patterns. These observations are consistent with selection having acted on entire networks of interacting loci. High-dimensional incompatibility networks can be viewed as multi-locus extensions of the classical DM model. While interesting from a data analysis standpoint, theoretical modeling of such multi-locus systems in the genomes of RILs is analytically not tractable, which makes this problem much less attractive from a theoretical standpoint. While the classical two-locus DM model represents a limiting case, it does give a plausible mechanistic description of how genetic incompatibilities initially arise in diverging sub-populations. Theory as well as empirical evidence suggest that, once DM-type incompatibilities take hold, additional incompatibilities accumulate exponentially (i.e. they “snowball”) (Orr and Turelli 2001; Matute *et al.* 2010; Moyle and Nakazato 2010). This exponential accumulation suggests that two-locus incompatibilities expand into multi-locus incompatibility networks over time, rather than accumulating independently in an additive manner.

In the present work we studied the selection signatures of different variants of the classical DM-model in genomes of RILs obtained by selfing. Our analysis showed that DM-type incompatibilities can give rise to complex inbreeding dynamics. In our view, the most troublesome situation is the presence of weak selection as it will continue to change genotype frequencies and LD patterns among RILs, even beyond genetic fixation. Hence, RILs that segregate incompatible genotypes do not, technically, present a reference population for the community, and phenotypic results may not be comparable across studies. Our analysis also showed that counter selection by breeders cannot rescue the adverse affects of genetic incompatibility but introduce additional biases in the form of LD and haplotype distortions. While these issues can be concerning to breeders whose aim it is to create these populations for downstream complex trait analysis, many existing RIL genotype datasets may present a largely unexplored resource for studying the basic principles underlying genetic incompatibilities. A deeper understanding of the molecular mechanisms that cause genetic incompatibilities in the genomes of RILs may provide valuable insights into the molecular mechanisms that drive speciation events in the wild.

## Acknowledgments

We thank M. Shojaei Arani for discussions and for his contribution on the preparation of some of the formulas. This work was supported by grants from the Netherlands Organization for Scientific Research (to F.J. and M.C.-T) and by a University of Groningen Rosalind Franklin Fellowship (to M.C.-T).

## Appendix

### A. General transition matrix

General form for the transition matrix *T* in the Markov chain, where *w* and *w′* represent the fitness of a genotype (Table 1):

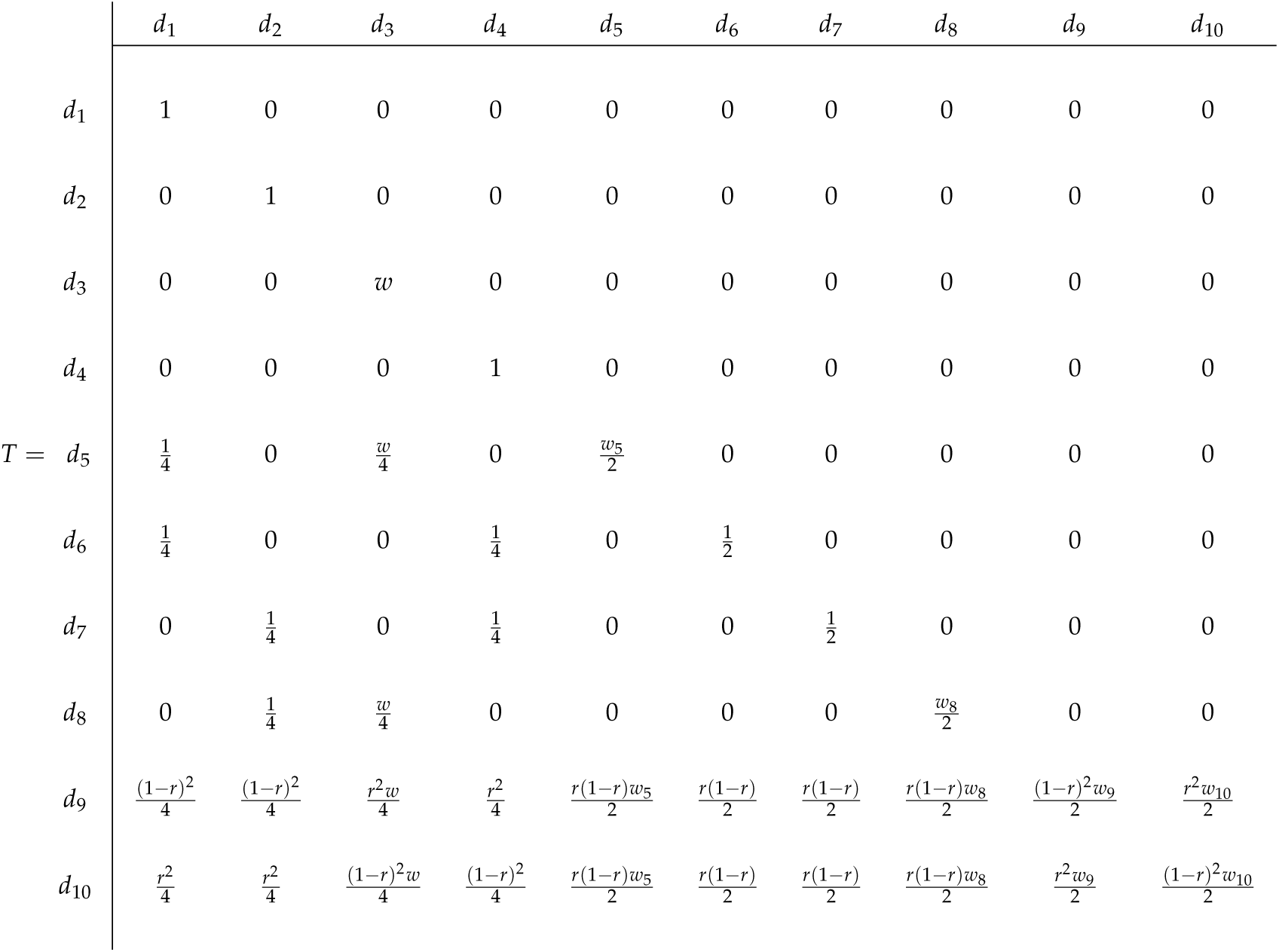

For *M*_1_, *w*_5_ = *w*_8_ = *w*_9_ = *w*_10_ = *w′*, for *M*_2_, *w*_5_ = *w*_8_ = *w*_9_ = *w*_10_ = 1 and for *M*_3_, *w*_5_ = *w*_9_ = *w*_10_ = 1 and *w*_8_ = *w′* (Table 2).

### B. Model M_0_ for F_∞_

The following equations show the diplotype and haplotype probabilities for the model without selection (*M*_0_) for *F*_∞_:

**Model M_0_ Diplotypes**

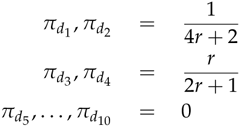

**Haplotypes**

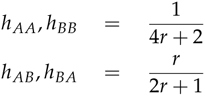

### C. Model M_0_ at intermediate inbreeding generations

The following equations show the diplotype and haplotype probabilities for the model without selection (*M*_0_) at intermediate inbreeding generations, where we used *u* = [(1 - 2*r* + 2*r*2)/2]*t* and *v* = [(1 - 2*r*)/2]*t*.

**Model M_0_ Diplotypes**

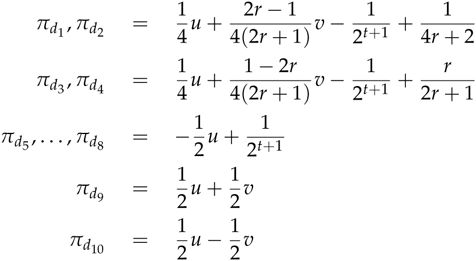

**Haplotypes**

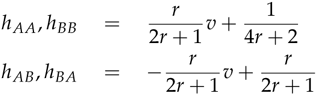

### *D. Models* M_1_, M_2_, M_3_ *for* F_∞_

Non-normalized diplotype and haplotype probabilities with selection for incompatibility models *M*_1_, *M*_2_ and *M*_3_ for *F*_∞_, for 0 *< w*, *w′ ≤* 1. The normalized expressions 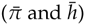 can be obtained dividing by 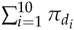.

**Diplotypes Model M_1_**

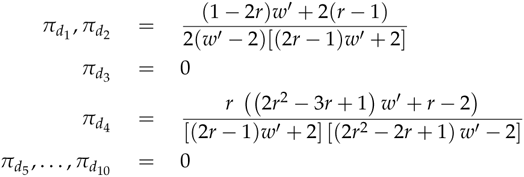

**Model M_2_**

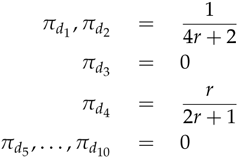

**Model M_3_**

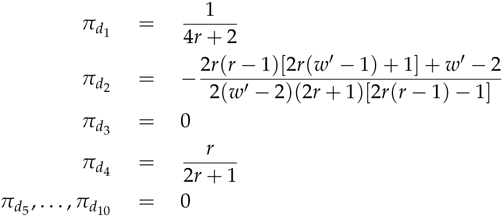

**Haplotypes Model M_1_**

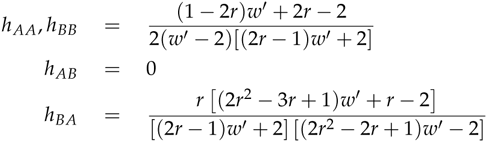

**Model M_2_**

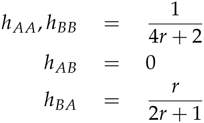

**Model M_3_**

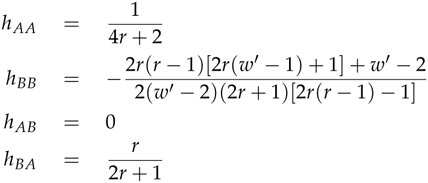

### E. Models M_1_, M_2_, M_3_ at intermediate inbreeding generations

Non-normalized diplotype and haplotype probabilities with selection for incompatibility models *M*_1_, *M*_2_ and *M*_3_ at intermediate inbreeding generations, for 0 *< w*, *w*^*′*^ *≤* 1. The normalized expressions 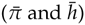 can be obtained dividing by 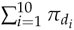. Note that *u* = [(1 - 2*r* + 2*r*^2^)/2]^*t*^, *v* = [(1 - 2*r*)/2]^*t*^, *u′* = [(1 - 2*r* + 2*r*2)*w′* /2]^*t*^, *v′* = [(1 - 2*r*)*w′* /2]^*t*^, *a* = 2*w -* (1 - 2*r*)*w′* and *b* = 2*w -* (1 - 2*r* + 2*r*2) *w′*.

**Diplotypes Model M_1_**

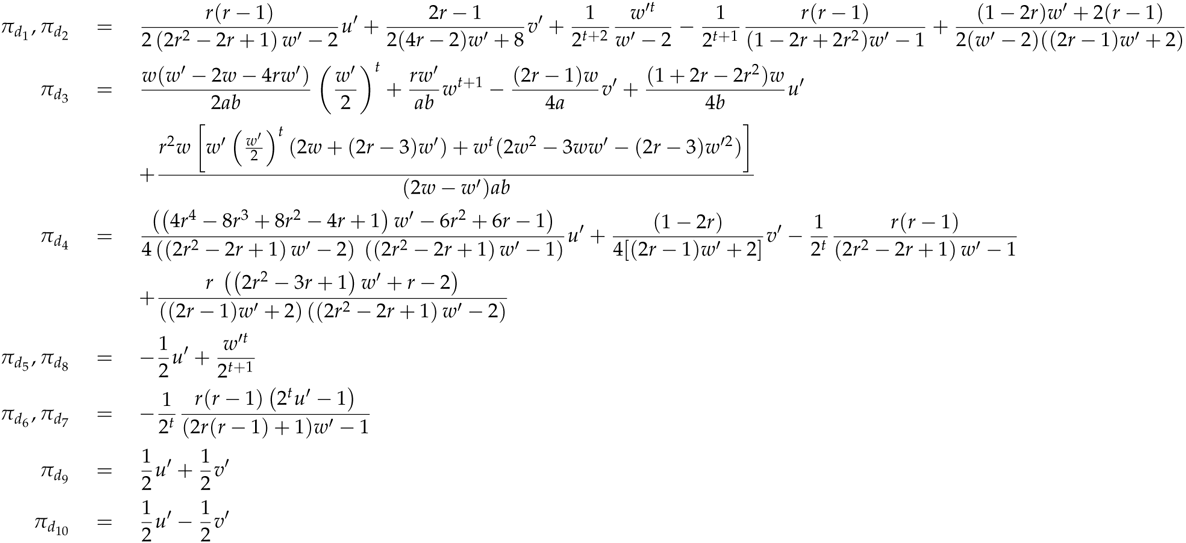

**Model M_2_**

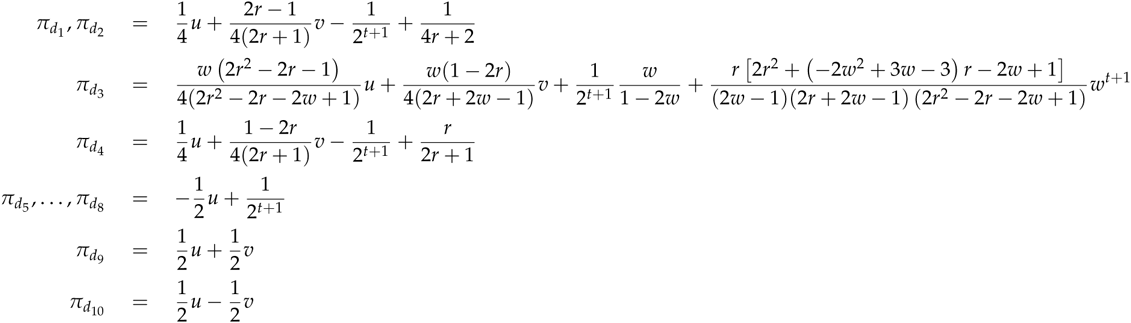

**Model M_3_**

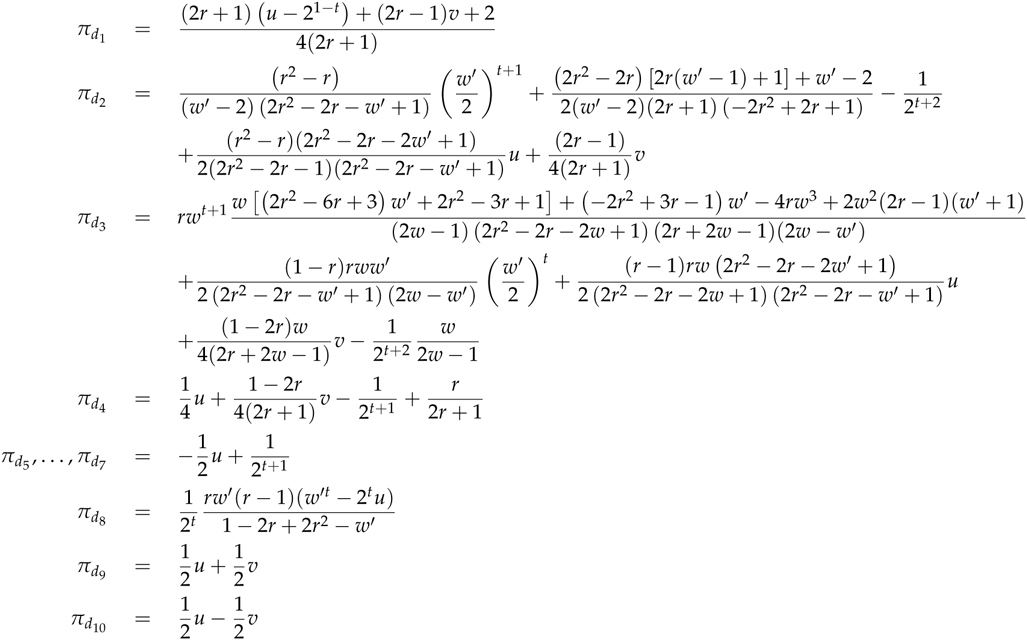

**Haplotypes Model M_1_**

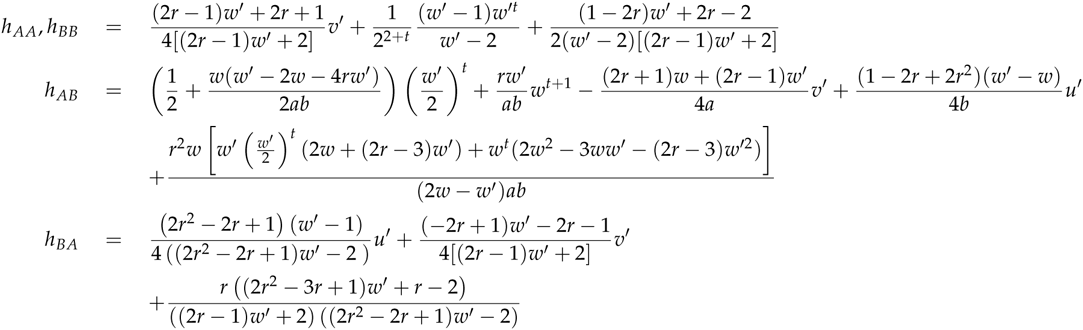

**Model M_2_**

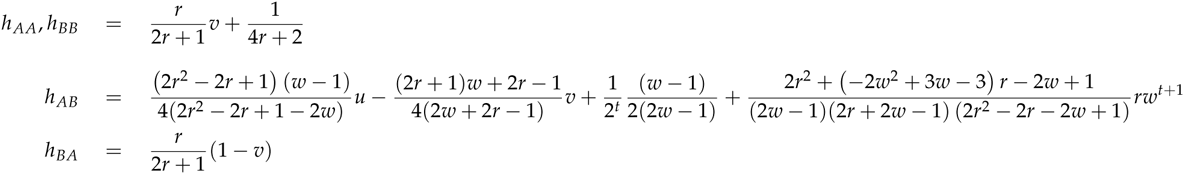

**Model M_3_**

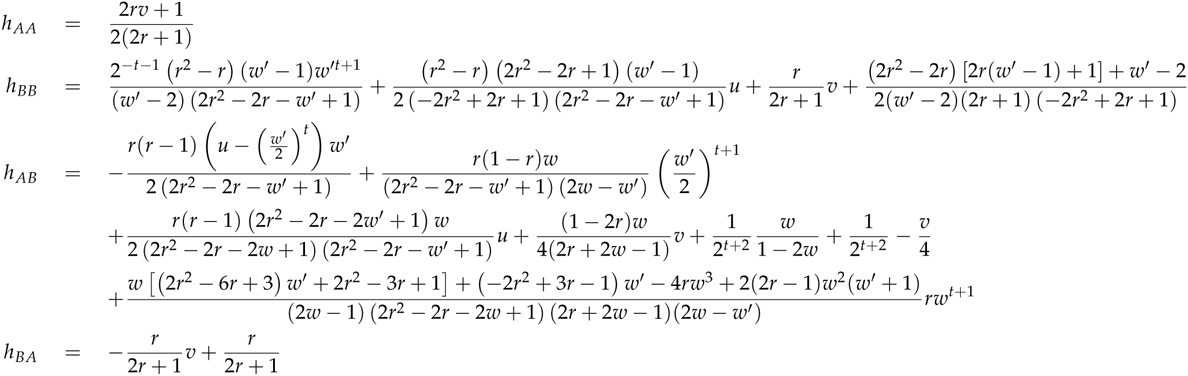

